# Multi-scale observations of mangrove blue carbon fluxes; the NASA Carbon Monitoring System BlueFlux field campaign

**DOI:** 10.1101/2022.09.27.509753

**Authors:** Benjamin Poulter, Frannie Adams, Cibele Amaral, Abigail Barenblitt, Anthony Campbell, Sean P. Charles, Rosa Maria Roman-Cuesta, Rocco D’Ascanio, Erin Delaria, Cheryl Doughty, Temilola Fatoyinbo, Jonathan Gewirtzman, Thomas F. Hanisco, Moshema Hull, S. Randy Kawa, Reem Hannun, David Lagomasino, Leslie Lait, Sparkle Malone, Paul Newman, Peter Raymond, Judith Rosentreter, Nathan Thomas, Glenn M. Wolfe, Lin Xiong, Qing Ying, Zhen Zhang

## Abstract

The BlueFlux field campaign is supported by NASA’s Carbon Monitoring System (CMS) and will develop prototype blue carbon products to inform coastal carbon management. Blue carbon is included in carbon-dioxide removal actions proposed to reduce atmospheric CO_2_ concentrations to mitigate climate change. Due to their high productivity and carbon storage, combined with historic losses and a wide-range of beneficial ecosystem services, the restoration and conservation of mangrove ecosystems features prominently in blue-carbon planning. The goal of BlueFlux is to carry out multi-scale measurements of CO_2_ and CH_4_ fluxes using chambers, flux towers, and aircraft and scale these to gridded products using space-based observations of forest structure and surface reflectance. The measurements cover gradients in disturbance, mainly from the history of hurricanes in the region that drive the dieback of mangroves and the formation of ‘ghost forests’. The fluxes of CH_4_ emissions will be contrasted with CO_2_ uptake to provide a more complete budget of radiative forcing and to understand the net climate benefits of blue carbon. BlueFlux demonstrates that quantifying the removals of CO_2_ and emissions of CH_4_ using a multi-scale approach can provide increased confidence in regional greenhouse-gas accounting, contribute to process-understanding, and help inform restoration and conservation efforts in the context of climate mitigation.

## 1. Introduction

Blue carbon is a key component in climate mitigation strategies that aim to reduce atmospheric carbon dioxide (CO_2_) concentrations through coastal and open-ocean carbon sequestration (Mcleod et al. 2011, Macreadie et al., 2019). At global scales, blue carbon forms part of the land-mitigation portfolio that could contribute to the Paris Agreement goal of keeping warming to well below 2.0-degrees Celsius and for achieving net-zero greenhouse-gas emissions (Roe et al., 2019). The high primary productivity of mangroves, salt marshes, and sea grasses, combined with restoration and conservation, is estimated to potentially store on order of an additional 1-5 PgCO_2_e yr^-1^ over present-day rates (Griscom et al., 2017). Given the wide range of services that coastal ecosystems provide, and their historical losses from land-use change (Goldberg et al., 2020), blue carbon could incentivize coastal restoration and protection through carbon financing (Zeng et al, 2021) and help avoid large losses of mangroves projected for the future (Adame et al., 2021).

Nature-based climate solutions aim to enhance both carbon uptake and ecosystem co-benefits (Seddon et al., 2022), yet there are still risks involved from both a social and scientific perspective (Macreadie et al, 2019). These risks are related to the permanence of carbon stored and potential displacement of land-use change elsewhere, i.e., leakage, and not knowing whether protection or restoration would have happened without carbon-based incentives, i.e., additionality. In addition, concerns about trade-offs between carbon-based management and biodiversity, water resources, food security and energy balance need thorough investigation (IPCC 2019). Unique to blue carbon is that coastal ecosystems also emit methane due to the presence of anoxic soils and methanogenic bacteria (mostly archaea). Methane (CH_4_) is a potent greenhouse gas with a 100-year global warming potential (GWP-100) ∼28 times greater than CO_2_ (Forster et al., 2021) and thus the climate mitigation potential of wetlands must be assessed by considering both carbon dioxide (CO_2_) removals and CH_4_ emissions (Rosentreter et al., 2018b). Measurements of these two trace gases, however, are sparse, with observations from soil chambers and flux towers limited to small geographic regions or over short time periods, leading to regional and global budgets that are highly uncertain (Rosentreter et al., 2021).

Mangrove ecosystems are of particular interest from a blue carbon perspective as they are one of the most productive ecosystems on Earth, with net primary production (NPP) ranging from 1000-2000 gC m^-2^ yr^-1^ (Alongi 2020). While only covering a fraction of the Earth’s land surface, 140,260 km^2^ (Bunting et al. 2022), they contribute ∼210 TgC yr^-1^ to global NPP (Alongi 2014). Much of this carbon becomes stored in biomass or sequestered in soil sediments, with recent lidar and radar estimates of total mangrove carbon stocks estimated around 5.03 PgC (Simard et al. 2019; & references within ranging from 1.32 to 11.2 PgC). These carbon stocks are concentrated in just a few key biogeographic regions, e.g., ten countries account for over 70% of total carbon stocks (Simard et al., 2019), which means that at national scales, mangrove carbon management can play a large role in nationally determined contributions and climate mitigation.

In contrast to CO_2_, global fluxes of methane from mangrove systems range from 0.2 to 1.5 Tg CH_4_ yr^-1^ (Rosentreter et al., 2021), a relatively small fraction of total wetland methane emissions (180 to 431 Tg CH_4_ yr^-1^, Rosentreter et al., 2021, Saunois et al., 2020). However, expressed in CO_2_-equivalents using GWP-20 and -100, Rosentreter et al., (2018) estimated methane emissions of 6.14 and 2.53 Tg CH_4_-Ce offset about 10-20% of global mangrove carbon sequestration (31.3 TgC yr^-1^). Considerable variability in annual mangrove methane fluxes due to climate, species, disturbance history, and salinity, and from uncertainties due to methodologies affect how well we know the ratio of carbon uptake to methane efflux. For example, based on scaling from flux-tower in mangrove systems, Delwiche et al. (2021) estimated 1.1 to 1.2 gCH_4_ m^-2^ yr^-1^, and from chambers, Rosentreter et al., (2018b) estimated 1.0 to 1.9 gCH_4_ m^-2^ yr^-1^, or 339±106 μmol m^−2^ d^−1^. Recent work using flux-tower observations tends to show that mangrove methane fluxes are slightly higher than the average of global annual wetland CH_4_ fluxes 1.1±0.2 gCH_4_ m^-2^ yr^-1^ (Knox et al., 2019), which would suggest the offset of warming from carbon sequestration is lower than previously estimated.

In 2020, the NASA Carbon Monitoring System provided support to establish the BlueFlux field campaign with the objective to develop prototype CO_2_ and CH_4_ products to inform mangrove restoration and conservation. The BlueFlux field campaign is designed to provide comprehensive measurements of CO_2_ and CH_4_ fluxes across Southern Florida and the Caribbean, with a focus on mangrove systems, their seasonal dynamics, and the adjacent ecosystems such as extensive sawgrass marshes and tree ‘islands’ within. This paper describes the BlueFlux multi-scale approach being implemented to measure fluxes from the water surface and the soil-root-stem to atmosphere pathways using chambers, and from ecosystem to atmosphere via aircraft eddy covariance and flux tower facilities. These flux measurements cover a gradient of ‘healthy’ mangroves, to recently disturbed and dying mangrove ‘ghost forests’ to help understand any directional change in carbon fluxes from losses and recovery (Figure 1). The flux measurements will form the basis of training data for machine learning algorithms and space-based observations to develop a daily, 500-m gridded surface flux product for years 2001-2022. The BlueFlux field campaign will contribute to quantifying how blue carbon can contribute to climate mitigation as well as contribute to reducing the uncertainty of temporal and spatial components of the mangrove carbon cycle.

**Figure 1:**
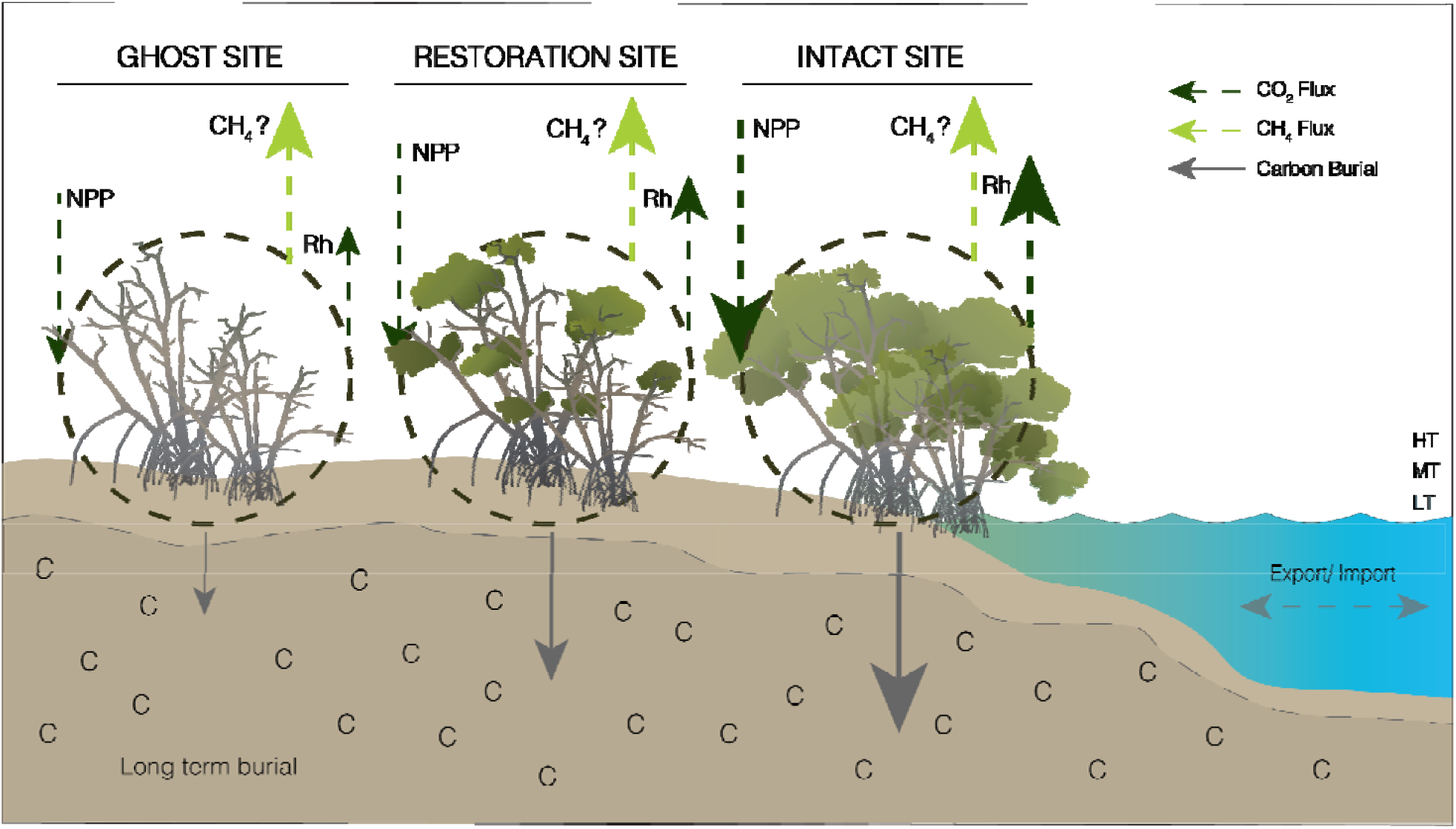
Schematic overview of BlueFlux’s objectives to compare and contrast net carbon uptake (NPP, burial), heterotrophic respiration (Rh) producing CO_2_ with methane greenhouse-gas (GHG) release across the existing disturbance and recovery gradients in Southern Florida.

## 2. The Region: Blue Carbon in Southern Florida and the Caribbean

BlueFlux is working within the Caribbean and Mesoamerican region, which includes the Gulf of Mexico, the islands of the Caribbean, and the coastal zones of Central and parts of South America (Figure 2). The coastal ecosystems within this region have been extensively impacted by development, hurricanes that cause erosion, storm surge and mangrove dieback (Sippo et al., 2018, Taillie et al. 2020; Lagomasino et al. 2021), sea-level rise (Parkinson and Wdowinski, 2022), and freeze events at the northern range limits (Cavanaugh et al., 2014, Osland et al., 2019). The losses of mangroves in the region are mainly driven by erosion and human-related activities (Goldberg et al., 2020) and are already expected to have foregone carbon sequestration opportunities in the range of 100s Tg CO_2_e yr^-1^ (Adame et al., 2021).

**Figure 2:**
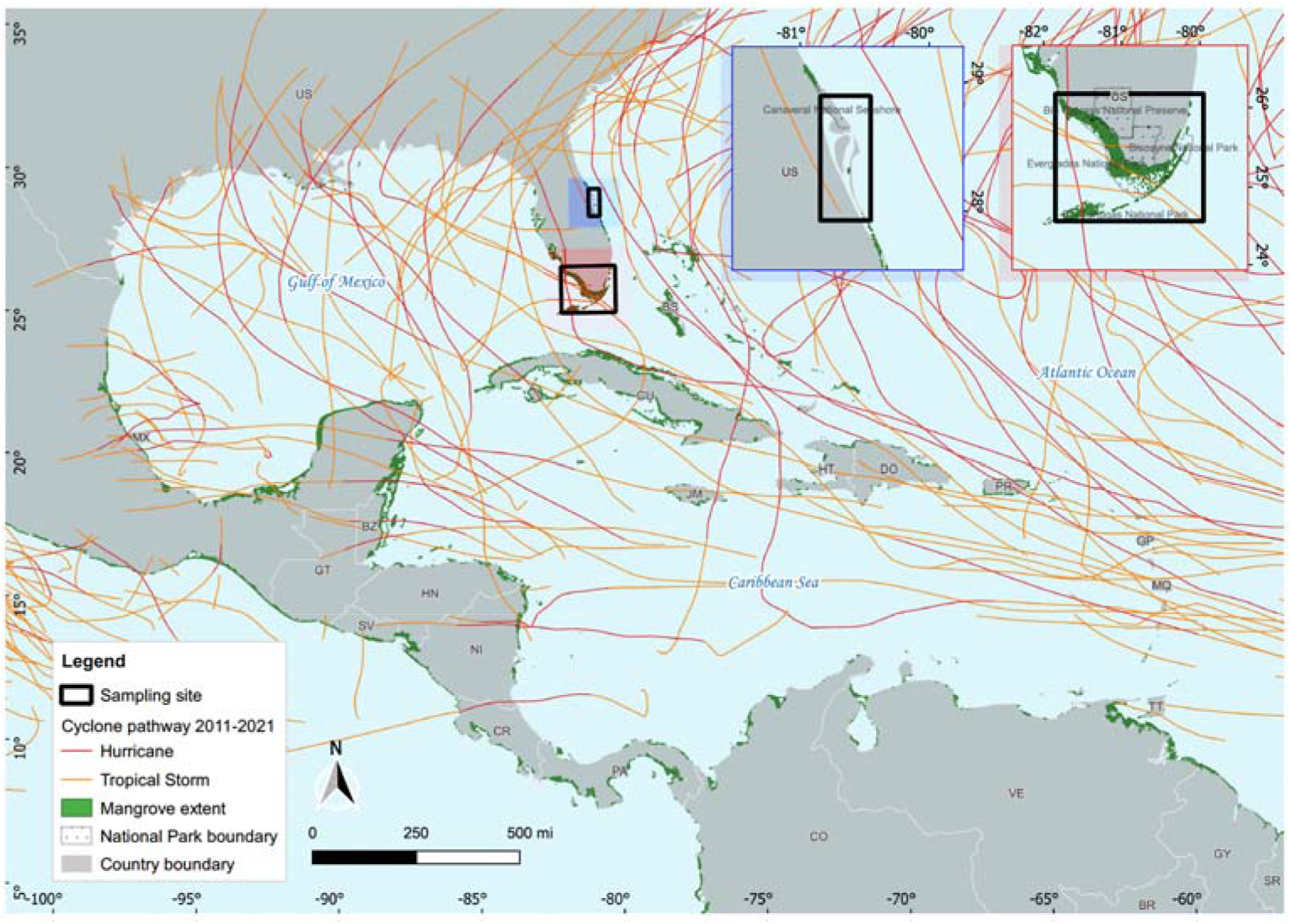
The Gulf of Mexico study region, showing mangrove extent (green) and the paths of tropical storms and hurricanes from 2011-2021 that drive the losses of mangroves through erosion and wind damage. Insets show the areas where the airborne component of BlueFlux will carry out eddy covariance measurements.

Three mangrove species grow in the Gulf of Mexico; red mangrove (*Rhizophora mangle*) with their distinctive aerial prop roots; black mangrove (*Avicennia germinans*) that have pneumatophores providing stability and oxygen to roots; and white mangroves (*Laguncularia racemosa*) with a smaller-glands at the leaf base to excrete salt. The distribution of each species is controlled by several factors including geomorphology, tidal connectivity, and soil properties (Snedaker, 1982). These factors can also influence the production of trace-gasses which measurements made in BlueFlux will aim to characterize across all mangrove species and their surrounding soils. Scaling this information will extend individual chamber measurements to the forest stand and to the landscape to better understand how gradients in species composition, salinity, and stature explain variability in carbon fluxes.

In-situ ground and aircraft measurements target areas within the conterminous USA, and ecosystems in Southern Florida that include a mix of ownerships (i.e., private, state, tribal and Federal). The Everglades and Big Cypress National Parks, which have a long history of biogeochemical research in association with the Florida Coastal Everglades Long-term Ecological Research Network (supported by the National Science Foundation), provides linkages with historical and contemporary research activities. Several flux towers within the Park boundaries provide information for both carbon dioxide and methane exchange and data from flux towers in Panama (managed by the Technological University of Panama in the Juan Diaz mangroves) and one in the Yucatan (at Puerto Morelos, Quintana Roo, Alvarado-Barrientos et al., 2021) will be used for sites outside the United States.

Within the Southern Florida core region, carbon stock and flux measurements will be made to understand how species, disturbance, hydrologic and climatic gradients explain flux variation. Six field campaigns, consisting of ground-based and airborne-based measurements are planned from 2022-2024, with the first field campaign successfully carried out in March and April of 2022. The following sections describe the field campaign from the perspective of 1) ground measurements being made for structure, species and trait-variation and chamber fluxes, 2) the tower flux measurements, 3) the aircraft measurements, and 4) the data-driven upscaling approaches using satellite remote sensing.

### 3.1.1 Ground measurements: mangrove forest inventory

Mangrove forest inventory data were collected at locations ranging in condition, with a primary focus on recently disturbed, dead, or regenerating mangrove ghost forests caused by recent extreme weather events. This information will be used to estimate plot-level forest wood volume and standing and dead biomass density. At each plot location a 10 m x 10 m plot was created, with the center of the plot oriented in the north and south cardinal directions. A GPS location was recorded at the plot center and at each plot corner using a WGS84 reference coordinate system. All trees taller than 0.5 m were tagged with a unique ID and the diameter and height of the tree were recorded using a Diameter-at-Chest-Height (DBH) tape and a hypsometer, respectively. The species and condition (i.e., live or relative decomposition of dead) of each tree was also noted. For plots with seedlings, we identified the species and measured height and basal diameter of each seedling in a 1 m x 1 m subplot. To record the quantity of debris material on the forest floor five 10 m transects were laid out every 72° starting along the North (0°) cardinal direction. At every 1-m interval along each transect, the size of any fallen woody debris that intersected the line was recorded. Within each of the 1 m intervals, a tally of the size of any fallen woody debris was noted. Woody debris size was categorized based on its diameter as fine (< 2 cm), small (2 cm), medium (4 cm), large (6 cm), and extra-large (> 10 cm).

### 3.1.2 Ground measurements for mangrove forest structure and volume

Ground measurements of three-dimensional (3D) mangrove forest structure and volume will be made using a terrestrial laser scanning system to supplement field plot based forest inventory data. These measurements will provide information on stem density, vertical distributions of biomass, and stand volume and surface area. A terrestrial scanning RIEGL VZ-400i instrument and two Global Navigation Satellite System (GNSS) units will be used to collect lidar point cloud data in the field (Figure 3). The integrated GNSS-TLS system has been optimized in the lab and through prior studies and its performance is deemed to be robust for coastal systems (Zhou et al. 2017; Xiong et al. 2017; Xiong et al. 2019). The scanner is programmed to measure distances by emitting a high frequency of laser pulses and capturing the return pulses when they are reflected from the surfaces of objects. The wavelength of the laser pulses is 1550 nm and the pulse rate can be up to 1200 kHz. The range measurement for the scanner can be as far as 880 m with an accuracy of 5 mm. The field of view is 360° in the horizontal plane and 100° in the vertical plane.

**Figure 3:**
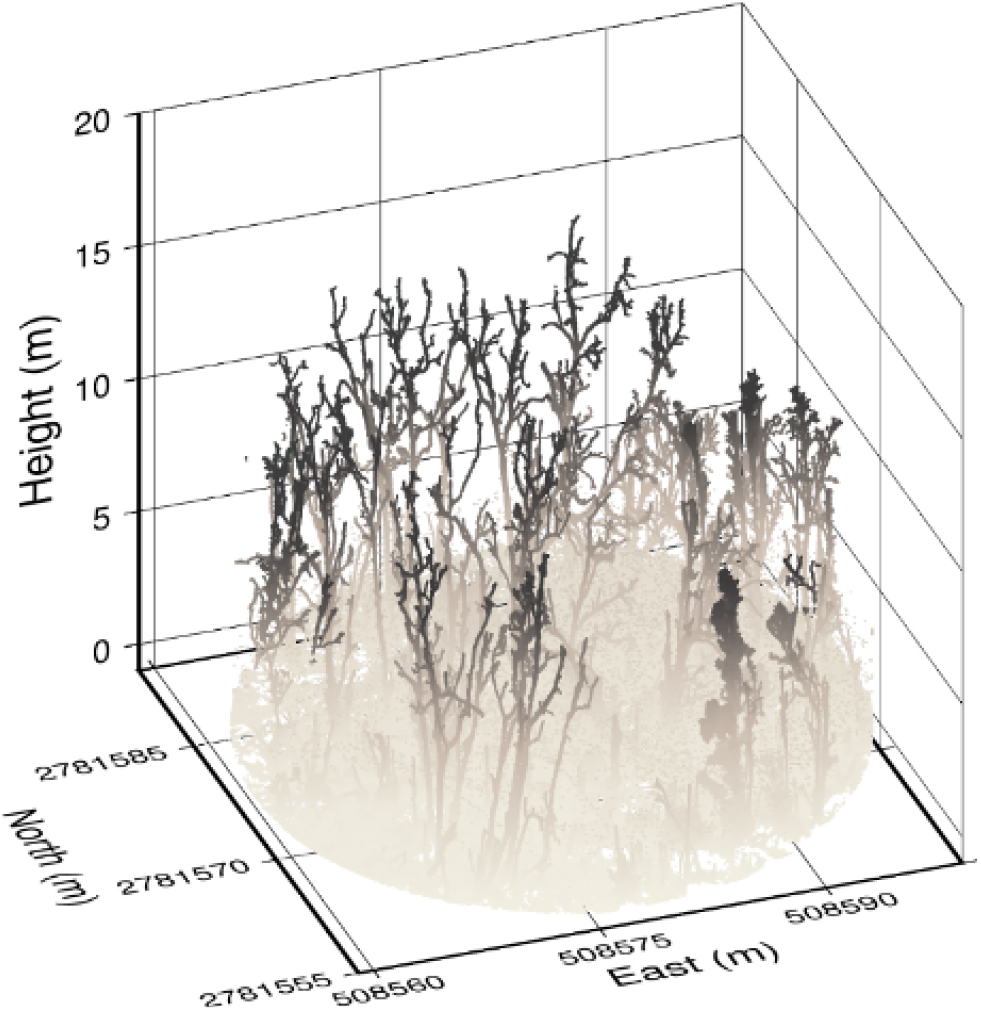
Example of a terrestrial laser scan of the mangrove system at Flamingo, Everglades National Park. The scan provides information on mangrove structure, including density and vertical profiles, as well as volume information that can be used to estimate biomass.

In each field plot, four panorama scans are collected around the center of the plot to reduce occlusion effects from branches and stems. Each scan takes ∼47 seconds and provides about 24 million points. The scan resolution will be set to be 0.03 degree in both horizontal and vertical directions. The GNSS unit mounted on the scanner was Trimble R8 as a rover station. The other GNSS unit, a Trimble R6, will be set up as a base station near the field plot. The observation time for the base station will exceed 4 hours each day providing centimeter-scale accuracy. During each scan, the position of the scanner will be measured by R8 with Real-Time Kinematic (RTK) method. In post-processing, point clouds from each scan will be registered in a local ‘project’ coordinate system by automatic data registration method in RISCAN PRO software. The core methodology is to apply an Iterative Closest Point (ICP) method in overlapped areas between multiple scans (Chen and Medioni, 1992). The registered point cloud will be georeferenced with the GNSS coordinates of four scan positions. Standard errors for registration and georeferencing are expected to be under 1 cm. Final point clouds of each field plot were outputted in ‘las’ format. The horizontal coordinates are WGS 84 / UTM zone 17N and the vertical datum is EGM96 geoid. We expect to make on order of 50-100 scans in a mix of mangrove stands (from healthy to dead or dying) based on landscape position and disturbance history.

### 3.1.3 Ground measurements: mangrove species composition

Spectral reflectances of vegetation provides biophysical information that can be used to understand and infer species composition, vegetation health, phenology, and sub-canopy water and soil properties. Leaf-level measurements of the spectral reflectance of the main plant species and soil and water surfaces across Southern Florida will be archived in a local database. The Analytical Spectral Devices, Inc. (ASD) spectroradiometer was used to acquire visible, near-infrared, and short-wave infrared spectra ranging from 350 to 2500 nm at 10 nm resolution, through a fiber cable. The absolute reflectances of classes of trees, grasses, shrubs, soil, and water were calculated with target object measurements divided by a simultaneous white panel measurement. The sensor probe uses a 25-degree view angle, at about 30 cm distance from the target object, to collect sample spectra at selected sites of vegetation and soil within ‘ghost’ mangrove stands, healthy mangroves with gaps, inundated grass, or shrubs. Reflectance samples were measured at four different positions from the sun, one on top (nadir) and three at different sun angles. For each position, the device recorded the average of 25 discrete measurements. Each reflectance measurement was normalized for incoming solar radiation using the reflectance of a white Spectralon panel. All the measurements were taken under a clear sky or with as few clouds as possible. The absolute reflectance of a sample was calculated by averaging the spectral values taken at the four sun angle positions of an object. Plots of selected samples and corresponding reflectances are shown in Figure 4.

**Fig 4.**
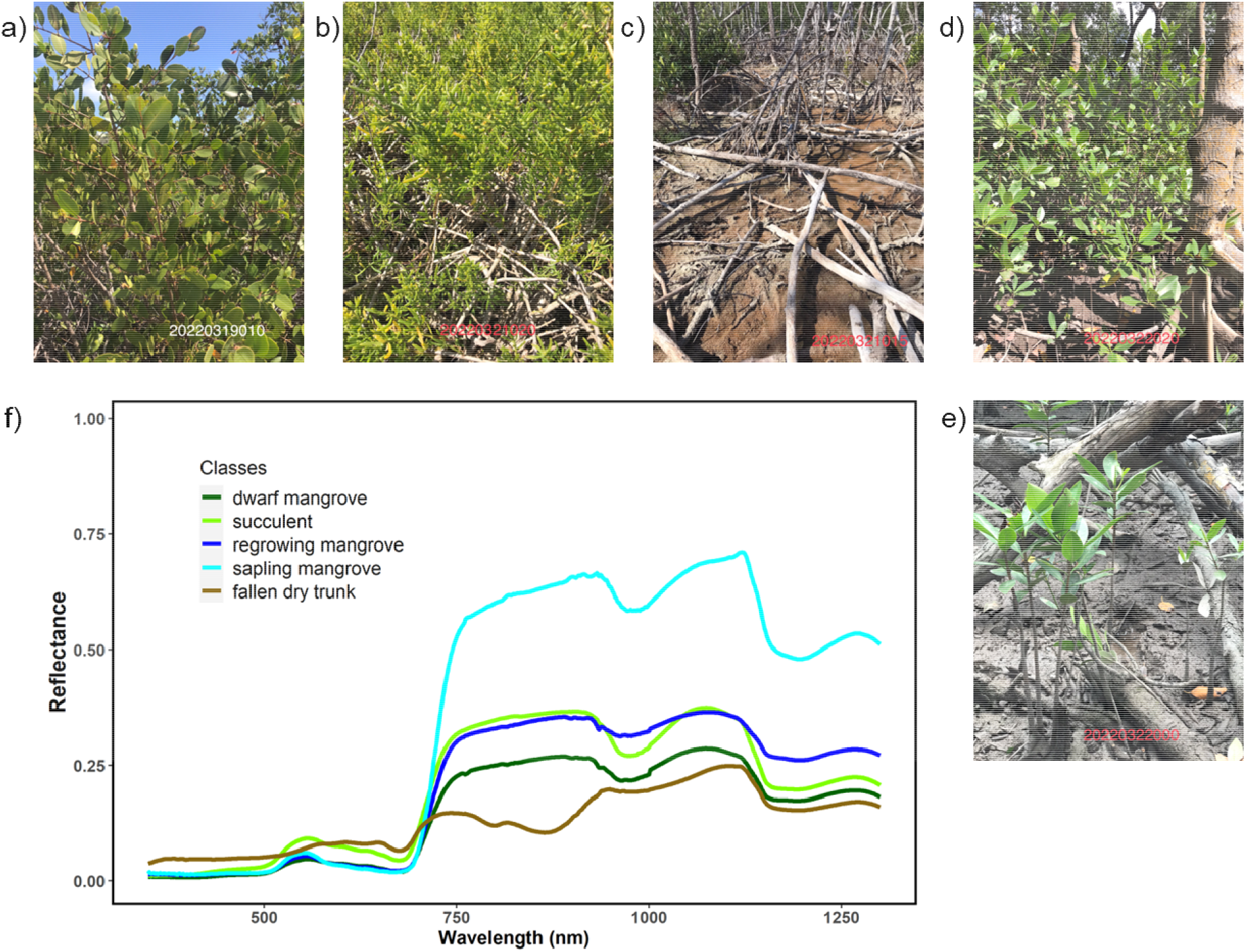
Spectral reflectance curves of the field-measured data: field pictures of (a) a dwarf mangrove, (b) succulent, (c) fallen dry trunks of dead mangroves, (d) regrowing young mangroves in a gap of a healthy mangrove forest, (e) mangrove saplings in the gap, and (f) corresponding reflectances measured by ASD spectrometer.

### 3.1.4 Ground measurements: soil and vegetation fluxes with chambers

The methods for estimating greenhouse-gas fluxes have been relatively well-established for quantifying soil, water, root, and stem exchange with the atmosphere. However, the relative contribution of these components to the overall ecosystem flux is still an active area of research. For example, recent studies have shown that gas fluxes from tree stems can be equally as relevant to the total ecosystem budget (Pangala et al., 2013, 2015, 2017). And a study by Jeffrey et al., (2019) found that methane fluxes from Australian mangroves themselves are also considerable relative to the overall ecosystem budget. In 2022, researchers published similar data that provided the first evidence that the aerial pneumatophore roots of mangroves, specifically for *Avicennia marina*, released 84% of the total ecosystem methane (Zhang et al., 2022).

In the first BlueFlux field campaign, tree, root, and soil CO_2_ and CH_4_ fluxes were measured in March 2022 at two highly degraded and two intact/regenerating forest sites within Everglades National Park (Figure 5). Fluxes were measured within plots previously scanned for forest structure and volume (Section 3.1.2) to allow for scaling of areal fluxes from ecosystem components to stand-level totals. Hand-made plastic chambers fitted with an inlet, outlet and a vent were used to measure the change of methane concentration over time from different plant and soil components. The Ultraportable Los Gatos Research Methane Analyzer and the portable Picarro GasScouter G4301 Mobile Gas Concentration Analyzer backpack were used to measure 1 Hz high-resolution CH_4_ and CO_2_ concentrations. Various chamber shapes were assembled and employed to account for the variety of shapes and sizes of the plant and sediment components (Figure 5; Troxler et al. 2015; Zhang et al. 2022), with the volume of each chamber calculated empirically by dilution of a methane standard (Siegenthaler et al. 2016; Jeffrey et al. 2020). These chambers included a ventilation hole to reduce pressure upon the surface when pushing and securing the chamber to the tree stem, soil, and root surfaces. This was plugged up with a rubber stopper after the chamber was affixed to the tree. Stem and prop root chambers were sealed directly to the plant surface using potting clay (Amaco). Pneumatophore chambers enclosed a single pneumatophore, with neoprene foam and potting clay used to seal around the base. Soil chambers were sealed, using vacuum grease (Dow Corning), to 8 inch diameter PVC collars, which were previously inserted 2 cm into the sediment at least 1 hour prior to sampling in order to avoid interference from sediment degassing. Chambers were tested for leaks using CO_2_ readings prior to each measurement.

**Figure 5:**
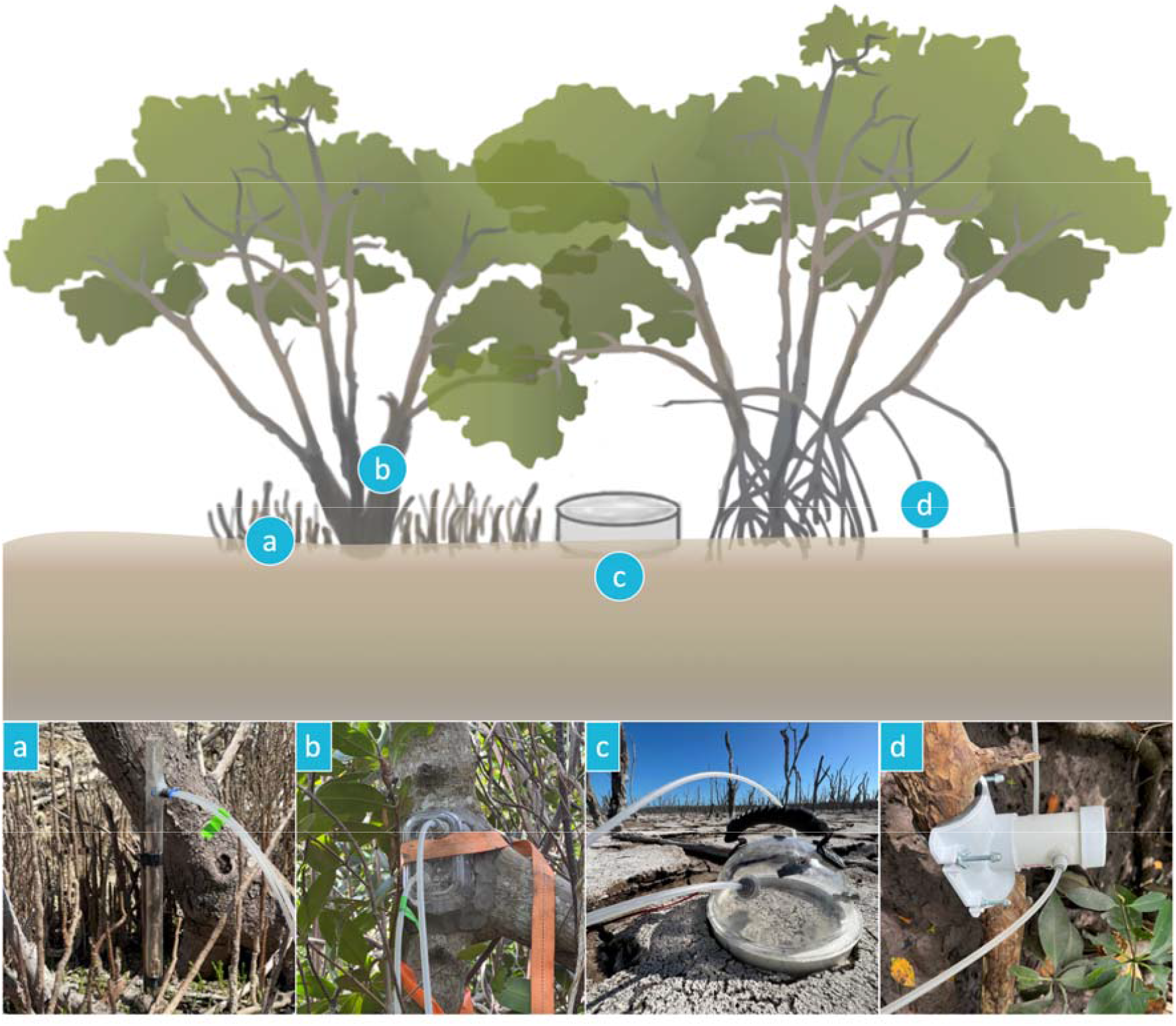
Diagram of ecosystem component fluxes measured via static chamber method and photographs of corresponding static chamber designs used: (a) pneumatophores, (b) mangrove stems, (c) sediments, (d) prop roots.

At each site, for sediment measurements, 10 collars were distributed randomly around the plot. Further, a minimum of 5 standing stems were measured, for both living trees and snags. Stem fluxes were measured at 0, 50, and 100cm from the base of the tree in order to test for vertical gradients in stem efflux, which can be indicative of soil rather vs. stem origin of stem-emitted gasses (Barba et al. 2018). For red mangrove stems, an additional flux was measured at the base of a live prop root. For black mangrove stems, and additional flux was measured on an adjacent pneumatophore. For each stem, a variety of ancillary data were also collected, including diameter at breast height (DBH), perimeter at each measurement height, tree stem, soil, and air temperature. Chamber incubations occurred for a minimum of 2 minutes for soils and 3 minutes for plant components.

### 3.1.5 Ground measurements: water chemistry

To capture water-air GHG exchange and its variability in mangrove waters in the Florida Everglades, a three-day spatial survey (March 2022) was conducted by driving a houseboat from Coot Bay (25.184921, -80.907329), up the Joe River to the Shark River to Tarpon Bay and back while measuring pH, water temperature, salinity, CO_2_ and CH_4_ and N_2_O concentrations, and CO_2_ and CH_4_ stable isotopes. Surface water was pumped continuously from ∼0.5 m water depth to the on-board setup consisting of a ‘showerhead’ equilibrator that was connected via a closed air loop to two gas analyzers, a Picarro G2201i and a Picarro G2308. Surface-water conductivity (EC), dissolved oxygen (DO), temperature, pH, and CDOM were measured every minute using a calibrated multi-parameter sonde (Eureka Water Probes). Filtered sterilized discrete samples for spectrophotometric pH, dissolved inorganic carbon (DIC) and total alkalinity (Talk) were collected periodically and analyzed in the laboratory (at Yale University).

### 3.1.6 Ground measurements: flux tower observations

The eddy covariance (EC) method was used to measure the exchange of trace gasses (CO_2_ and CH_4_) between wetland systems and the atmosphere throughout the Florida Everglades. Several flux towers exist in the region as part of the Florida Coastal Everglades Long Term Ecological Research network (FCE LTER) along the Shark River Slough (SRS) and Taylor Slough/Panhandle (Ts/Ph) hydrologic gradients. The EC towers are equipped with 3D sonic anemometers (model RS-50, Gill Co., Lymington, England) that measure wind speed and sonic temperature, and use an open path infrared CO_2_/H_2_O gas analyzer (LI-7500, LI-COR, Inc., Lincoln, Nebraska), and a CH_4_ analyzer (LI-7700, LI-COR Inc., Lincoln, Nebraska). Flux data are measured at 10 Hz. In addition to flux data, meteorological measurements included net radiation (CNR 1, Kipp and Zonen, Bohemia, New York) and incoming and reflected PAR (model LI-190SB, LI-COR, Inc., Lincoln, Nebraska), air temperature (*T*_*a*_) and humidity (HMP45C, Campbell Scientific, Inc., Logan, Utah) and wind speed and direction (model 05103 RM Young, Traverse City, Michigan).

The original mangrove forest tower, SRS6, was constructed in June of 2003 and the instruments are installed 27-m above the soil surface on a 30-m tower. While measurements of CO_2_ and H_2_O go back to 2004, CH_4_ measurements began in 2018. Air temperatures are measured at 20 m, 15 m, 11 m, 6 m, and 1.5 m above the ground surface using aspirated and shielded thermometers (107 temperature probes, Campbell Scientific, Inc.). Below ground soil heat flux (HFT 3.1, Campbell Scientific, Inc.) and soil temperature (105T, Campbell Scientific, Inc.) (*T*_*s*_) are measured at 5 cm, 10 cm, 20 cm, and 50 cm. Hydrologic data at this site are continuously monitored and recorded every 15 min at a station 30 m south of Shark River and 150 m west of the flux tower. Measurements included specific conductivity and temperature (600R water quality sampling sonde, YSI Inc., Yellow Springs, Ohio) of surface well water and water level (Waterlog H-333 shaft encoder, Design Analysis Associates, Logan, Utah). The detailed information on construction of base of tower and other supporting structure can be found in Barr et al. (2010).

### 3.2.1 Airborne eddy covariance fluxes; CArbon Atmospheric Flux Experiment (CARAFE)

Airborne eddy covariance (AEC) is a well-established technique for quantifying surface-atmosphere exchange of trace gasses and energy (Desjardins 1982). When combined with wavelet transforms (Wolfe et al. 2018), AEC can characterize spatial gradients in fluxes at model-relevant scales (1-100 km). Flux footprint modeling allows for evaluation of fluxes within the context of surface properties and modeled fluxes (Hannun et al. 2020; Vaughan et al. 2021). Such data is complementary to ground-based observations, which integrate over a relatively small area but may better constrain site-specific processes and temporal variability.

Blueflux AEC observations employ the CArbon Atmospheric Flux Experiment (CARAFE) payload on a Beechcraft King Air A90 flown by Dynamic Aviation and equipped with meteorological and trace gas sensors. An AIMMS-20 (Aventech) provides 10 Hz observations of 3D wind velocities, air temperature, aircraft position, and aircraft orientation (pitch/roll/yaw). This system includes a probe (mounted under the left wing) for meteorological measurements coupled with high-resolution differential GPS and inertial navigation systems. Similar systems have been utilized for airborne EC (Vaughan et al., 2021). Ambient air is sampled through a gas inlet mounted under the right wing and transferred through a teflon tube (in the wing) to two gas sensors in the cabin. A Picarro G2401-m provides 0.5 Hz measurements of CO_2_, CH_4_, H_2_O, and CO, while a Picarro G2311-f provides 10 Hz measurements of CO_2_, CH_4_, and H_2_O. The G2401-m contains specialized pressure control systems for airborne operation and thus serves as the accuracy standard for mixing ratios, while the G2311-f provides the fast time response needed for AEC. Dry CO_2_ and CH_4_ mixing ratios are calibrated in the laboratory against NOAA WMO compressed gas standards with a two-point calibration.

Figure 6 maps the flight tracks from the April 2022 field campaign. Each deployment consists of 4-6 flights (25 flight hours per deployment), with 5 deployments planned over 2022 and 2023. The flights are designed to focus on coastal mangrove vegetation in both South and East Florida but also include mapping over inland forests and wetlands, such as the extensive sawgrass marshes (*Cladium jamaicense*). Flight duration ranges from 2.5 to 4.5 hours. Typical altitude is 100-m above ground level, with occasional spirals to ascertain mixed layer depth and flux legs higher in the mixed layer (200 - 800 m) to determine vertical flux divergence and corrections. At an altitude of 100 m, we expect the flux footprints to be roughly 5000-m wide with 50% of the flux within 1000-m and 90% within 5000 m for typical surface wind speeds of 5 to 10 m s^-1^. The fluxes for CO_2_ ranged from 0 to -40 μmol CO_2_ m^-2^ s^-1^ and the fluxes for CH_4_ ranged from 0 to 200 μmol CH_4_ m^-2^ s^-1^ (Figure 7 a and b). In general, the methane fluxes appear to be higher for sawgrass and CO_2_ uptake greater for mangroves for the April field campaign and further flights will explore seasonal and interannual variability.

**Figure 6:**
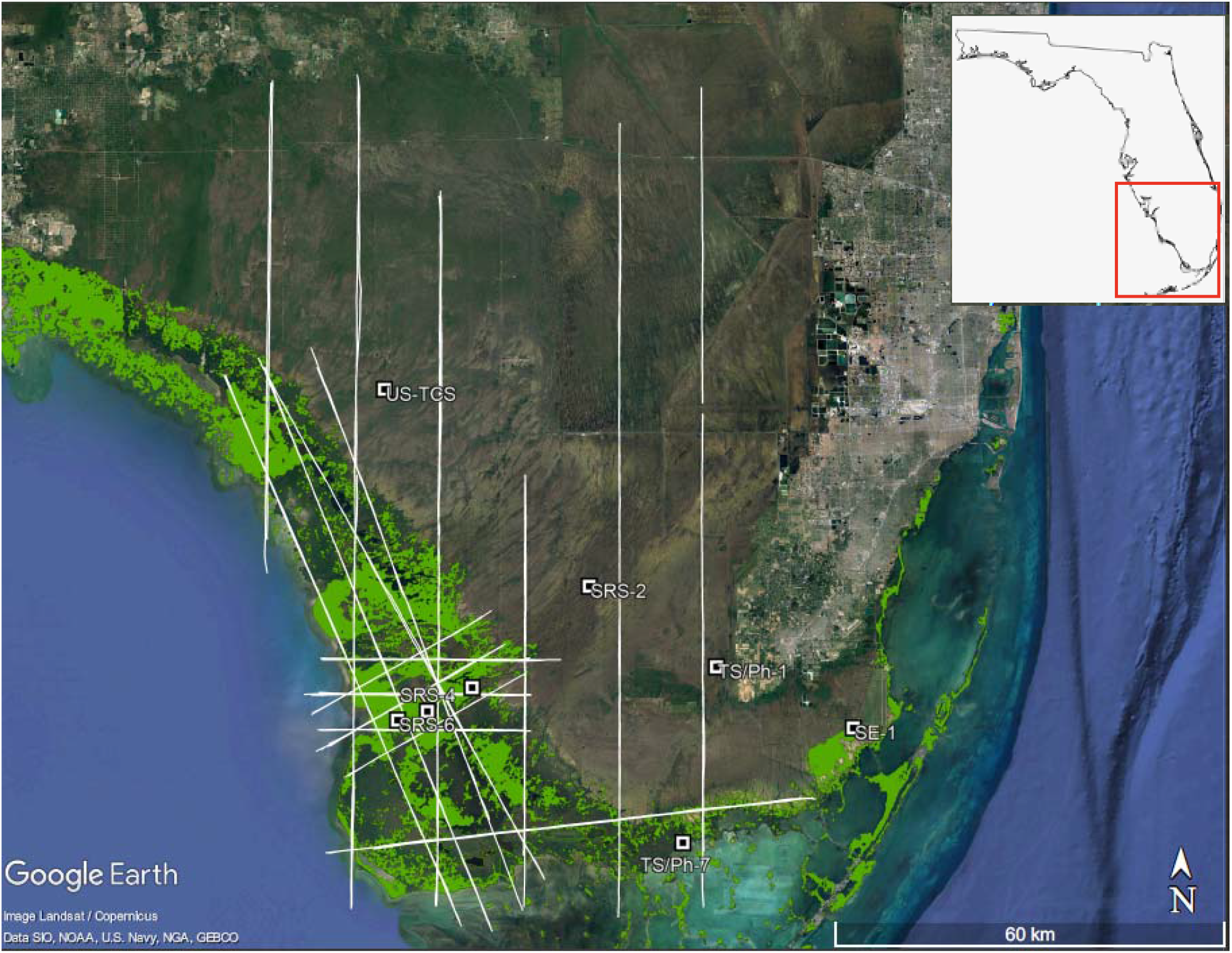
BlueFlux airborne operations in April 2022. White lines show tracks for airborne flux legs; note that each line may include multiple overlapping transects. White squares are long-term surface observation sites. Green patches denote Mangrove extent as of January 2022 (https://geodata.myfwc.com/datasets/myfwc::mangrove-habitat-in-florida-1/about, last accessed 21 June 2022) and the location of the SRS and TS flux towers are labeled. Inset show the location of the operation area in Southern FL.

**Figure 7:**
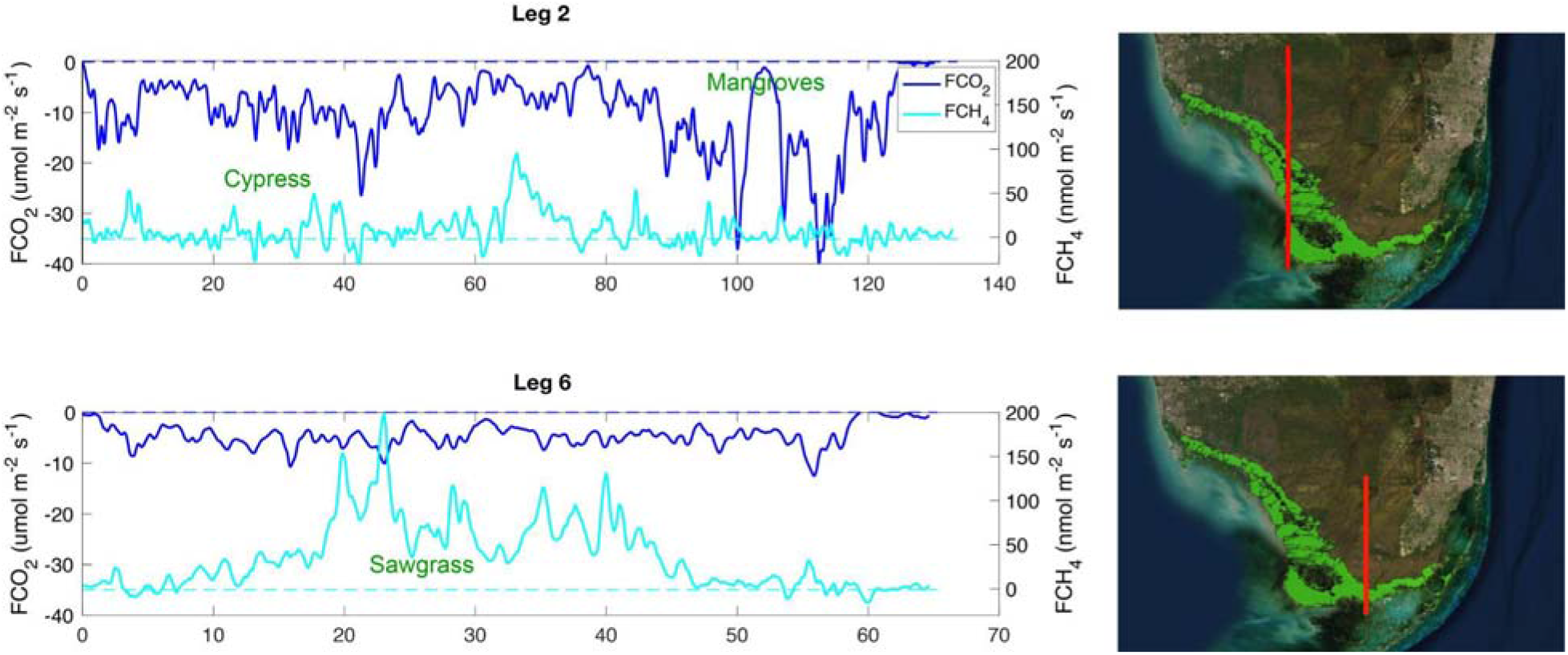
Time series of methane (cyan) and carbon dioxide (blue) fluxes on two legs of the Everglades raster flight from 21 April 2022. The x-axis is the distance traveled over each leg, with 0 denoting the northernmost point of each leg. Maps to the right of the time series depict the flight tracks of each leg in red. Green shading shows regions of mangrove habitats. The wind during this flight was from the north east at an average of 9 +/- 2 m/s.

NASA’s Commercial Air Service provides the basis for flight safety reviews. Authorization to proceed (ATP) is given after the flight planning follows NASA’s Airworthiness Review process (NPR 7900.32D) which initiates a series of technical interchange meetings (TIMs), flight test requirement documentation (FTRD), preliminary airworthiness review (PAR), final airworthiness review (FAR), and operational and operational/mission readiness reviews (ORR and MRR). The flight planning also follows the Section 7 Consultation to the Endangered Species Act and Section 106 of the National Historic Preservation Act (NHPA) in coordination with the tribal and state historical preservation offices (THPO and SHPO). After receiving permits from the National Parks, BlueFlux flights are coordinated each day with local airborne surveys and guided by near-real time meteorological information from the NASA Goddard Modeling and Assimilation Office Mission Tools Suite (MTS).

### 3.3 Prototype blue carbon products

The airborne flux measurements will be used as training data for a data-driven upscaled model that will provide daily gridded carbon fluxes. Given the spatial and temporal representation of the fluxes from the flights, the disaggregation of the ecosystem level fluxes to the components of water, soil and stem, and the validation with tower data, the gridded prototype products can provide the basis for a variety of studies. Figure 8 provides a preliminary example of the gridded methane fluxes using data from the seven flight days between April 19th and 26th, 2022. These flights provided measurements of methane fluxes for 43,141 sample points at 500-m resolution. We used the MODIS Nadir Bidirectional Reflectance Distribution Function (BRDF)-Adjusted Reflectance (NBAR, MCD43A4v006) data over the study area downloaded from Google Earth Engine (GEE). MODIS NBAR is developed daily at 500 m spatial resolution, using 16 days of Terra and Aqua data to remove view angle effects and temporally weighted to the ninth day as the best local solar noon reflectance (Schaaf et al. 2002; Wang et al. 2018).

**Figure 8:**
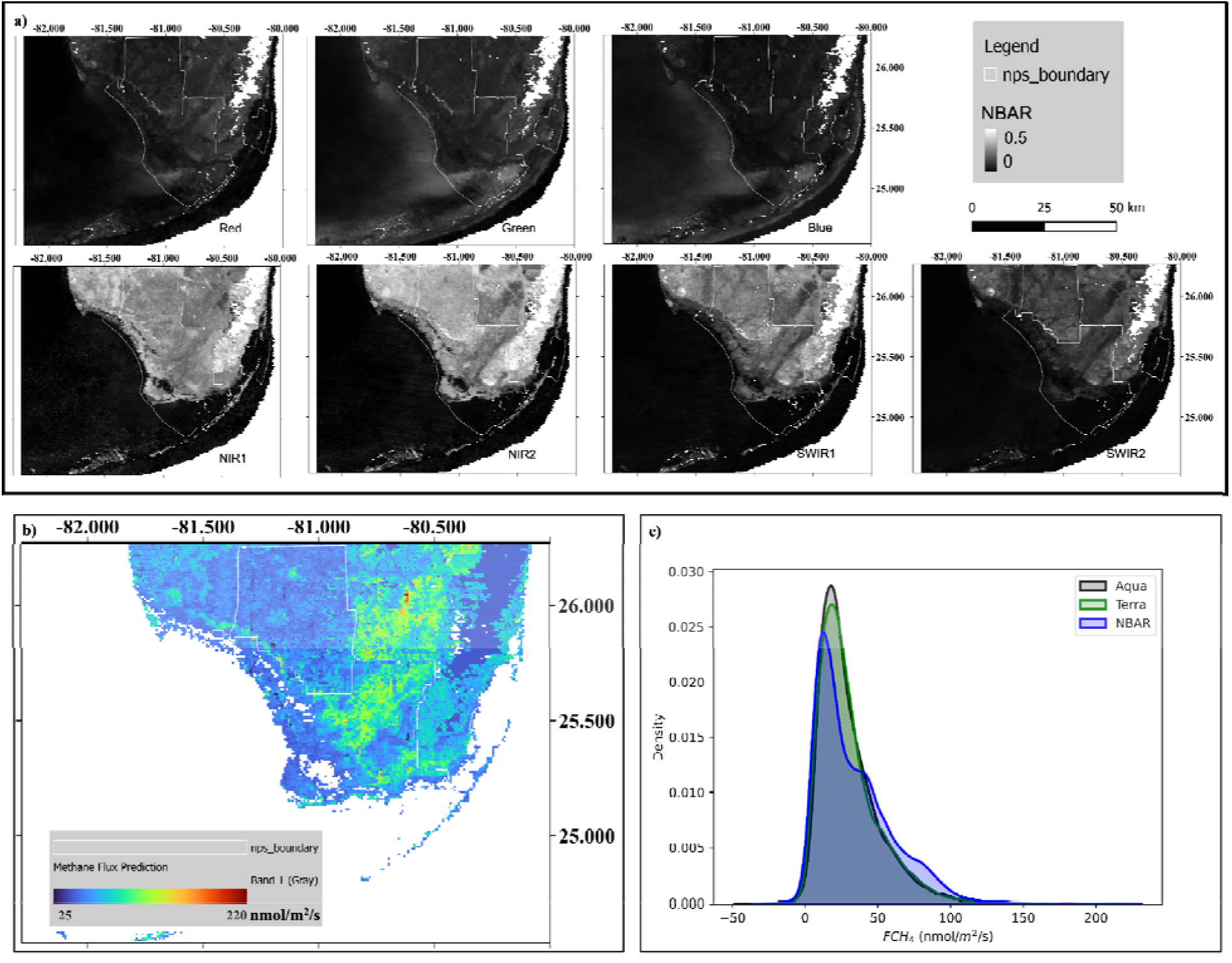
Data-upscaling for methane using flight tracks in Figure 6 to train a reflectance-based model using (a) MODIS nadir BRDF-adjusted reflectance (NBAR) of all seven land bands: red (620-670 nm), green (545-565 nm), blue (459-479 nm), NIR1 (841-876 nm), NIR2 (1230-1250 nm), SWIR1 (1628-1652 nm), SWIR2 (2105-2155 nm). The Everglades National Park boundary is shown in white polygons; (b) the gridded methane fluxes for April 2022; and (c) comparison between modeled fluxes from MODIS NBAR reflectance versus Terra and Aqua.

Mean values of MODIS NBAR product between April 19th and 26th were composited per band of seven spectral bands (Figure 8a, red, NIR1, blue, green, NIR2, SWIR1, SWIR2) as model input features. The MODIS Terra land-water mask (MOD44Wv006, Carroll et al. 2017) of 2015 was applied to mask open-water pixels. Then all the flight methane flux sample points at 5-km resolution were mapped to MODIS grids by averaging FCH_4_ values of points that fell in the same MODIS pixel. This produced 1528 gridded data. An upscaling model was trained at the MODIS pixel scale using ensemble random forest regressors (Breiman 2001; Kim et al. 2020). Random forest regression contains an assembly of independent trees constructed from a random subset of input data or input space (features). The generalization error converges as the forest grows to a limit, which avoids overfitting. We used the *scikit-learn* library in Python to build up random forest regressors with a bootstrap ensemble sampling for data. A ‘tree’ grew a split where a random selection of two features reduces the mean squared error (MSE) at a leaf node.

The model yielded an ‘out of bag error’ of 39.82 μmol m^2^ s^-1^. We applied the model to all the flight methane flux grids. The root mean squared error (RMSE) of model prediction is 10.66 μmol m^2^ s^-1^, and the coefficient of determination is 0.92. Red and near-infrared bands that have been widely used for characterizing vegetation chlorophyll, canopy structure and soil wetness were among the most important bands to model spatial variability of methane fluxes in the Everglades National Park area. We also trained separate upscaling models on MODIS Aqua and Terra 8-day composite level 3 surface reflectance data (MYD09A1v061 and MOD09A1v006, Vermote et al. 2002; Bréon et al. 2012). Only pixels with good quality data were used. The RMSE of Aqua model was 12.74 μmol m^2^ s^-1^ and 12.41 μmol m^2^ s^-1^ for Terra trained model. We applied the models on Aqua, Terra, and NBAR data to upscale FCH_4_ to the whole region. Methane fluxes mapped from NBAR data in our study area have a better spatial representation of methane emission gradients and variabilities than those from Aqua or Terra data (Figure 8c). High methane emissions were located in inundated sawgrass marsh in the Shark River Slough and the estuarine section of Taylor Slough (Figure 8b). Due to frequent cloud cover, the 8-day composite of Aqua or Terra data are noisier than the NBAR product over high emission marsh and low emission cypress swamp, which produced different density distributions of methane fluxes (Figure 8c).

#### Anticipated Results

One of the current shortfalls of ‘blue carbon’ assessments is that they consider carbon stores and stocks but often overlook non-CO_2_ greenhouse gas emissions that can greatly affect (positively or negatively) the overall net radiative forcing effect of these ecosystems. Mangroves are intertidal ecosystems and while net autotrophic at the ecosystem scale (Duarte et al. 2005, Alongi et al. 2014), creek waters and sediments are generally a source of atmospheric CO_2_ and CH_4_ (Rosentreter et al. 2018a; Call et al. 2015) and can also act as a source or sink for N_2_O (Maher et al. 2016; Reithmaier et al. 2020). Along the tidal elevation gradient (creek to forest basin), mangrove coverage, species diversity, and sediment structure can change markedly, resulting in great spatial variability of GHG fluxes. The inter-site variability of GHG fluxes from mangroves is further driven by various other factors including regional-climatic (e.g. rainfall, storms), hydrological (e.g. groundwater), geomorphological (e.g. bedrock, sediment composition, typology), physicochemical (e.g. salinity, temperature, oxygen), biotic (e.g. microbial community, bioturbation), biogeochemical (e.g. availability of organic matter and nutrients), and anthropogenic (e.g. agriculture, urbanization, pollution) factors (Rosentreter et al. 2021).

The BlueFlux field campaign is designed to collect detailed information on mangrove structure and multi-scale measurements of GHG fluxes. The data-upscaling approach described in section 3.3 will capture these edaphic, hydrologic and disturbance gradients either through the reflectance data or through additional covariates, such as wind speed, remote sensed forest structure using radar or lidar, and topographic information. The gridded carbon flux products will provide a basis for evaluating trends over the past two decades in GHG fluxes and their spatial patterns in response to changing climate and climate extremes, hurricane history, and land management.

#### Anticipated Impact

Blue carbon is integrated within many policy-guiding documents and climate mitigation policies themselves (Hilmi et al., 2021). The Intergovernmental Panel on Climate Change (IPCC) focuses on blue carbon specifically in the Special Report on the Ocean and Cryosphere in a Changing Climate (SROCC) and discusses the consequences of maintaining and restoring these ecosystems for climate mitigation. The Blue Carbon Policy Project of the International Union for the Conservation of Nature (IUCN) helped guide recommendations to the United Nations Framework Convention on Climate Change’s 26th Conference of Parties (COP26) in 2021. At COP26, blue carbon related goals included enhancing ambition, accelerating implementation and monitoring and verification of results.

A key aspect of NASA’s Carbon Monitoring System is to support stakeholder needs and expand partnerships with practitioners involved with climate mitigation. BlueFlux is partnered with stakeholders in 1) the private sector through the Everglades Foundation and the Environmental Leadership and Training Initiative (ELTI) at Yale University, 2) through federal agencies, including NASA, United States Geological Survey (USGS), the National Park Service, and National Science Foundation, 3) through tribal nations, including the Miccosukee, the Seminole Nation of Oklahoma and the Seminole Tribe of Florida, and 4) through international partnerships with the Coastal Biodiversity Resilience to increasing Extreme events in Central America (CORESCAM). The data access policy follows NASA’s Open Source Science philosophy of being freely and readily accessible, with transparent metadata and documentation.

BlueFlux intends to benefit these stakeholders by accounting for GHG exchanges from critical blue carbon ecosystems, i.e. mangroves and sawgrass marshes. Determining gas flux in sites under different pressures and conditions such as intact, degraded, or restored, and over time (intra- and interannual variability), is of ultimate need for informing national greenhouse gas inventories (NGHGI) more precisely. The multi-source, multi-scale quality of the project also aims to allow stakeholders to access information and data from the soil, and water to the atmosphere that may support the generation and refinement of Tier 3 models, and the assessment and reduction of the uncertainties associated with current models for NGHGI (Ogle et al., 2019). Cross-scale-calibrated flux estimates at 500 to 1000 m resolution which range in time and space across Mesoamerica and the Caribbean might also support local, and regional policies and projects on climate change adaptation and mitigation, ecosystem restoration, and carbon market. The project also provides data that can support the global stocktake of the Paris Agreement (GST) and the nationally determined contributions process, since it maps GHG flux time series from coastal ecosystems under varying conditions and, as consequence, highlights the history of regional carbon sinks and sources.

## Acknowledgements

The Blueflux project acknowledges core support from the NASA Carbon Monitoring System and NASA’s Terrestrial Ecology Program. We would like to thank the BNP-PARIBAS foundation for their support to the CORESCAM project (Coastal and Marine biodiversity resilience to increasing extreme events in Central America and the Caribbean), from their 2019 Biodiversity and Climate Change call. We also thank the Everglades and Big Cypress National Parks and the Florida Coastal Everglades Long Term Ecological Network for their support.

## Notes

### Competing Interest Statement

The authors have declared no competing interest.

